# Deep Learning for Cognitive Task Presence Prediction from Dynamic Functional Connectivity

**DOI:** 10.1101/2025.11.14.688561

**Authors:** Kini Chen, Mohammad Torabi, Jie Jian, Archer Y. Yang, Jean-Baptiste Poline

## Abstract

Dynamic functional connectivity (dFC) studies the time-varying coordination between brain regions measured with fMRI and is a potential biomarker for understanding cognitive dynamics and tracking the development of neurological disorders. However, a critical methodological challenge lies in the variability of dFC estimates across different dFC assessment methods, raising concerns about the reliability and interpretation of downstream findings. While deep learning (DL) models have demonstrated the ability to capture traditionally inaccessible data patterns in many disciplines, they encounter challenges when applied to neuroimaging data. For instance, the high dimensionality, noise, and temporal complexity inherent in dFC makes it challenging for DL models to extract meaningful and interpretable insights. In this study, we investigated how DL architectures can be developed and adapted to predict task presence over time from task-based dFC data, and additionally, how the choice of dFC assessment method influences the predictive performance of DL models. We developed and compared a convolutional neural network (CNN), a node-level classification graph convolutional network (GCN), and a graph-level classification GCN based on their ability to predict time points at which subjects were performing a cognitive task or at rest. In our study, the results indicate that both DL model architecture and dFC estimation methodology significantly impact task presence prediction capacity, while the specific task paradigm had minimal influence out of the limited types that were explored. This work offers a powerful benchmark for understanding the dynamics of underlying task-driven cognitive state transitions and the analytical flexibility limitations of dFC estimation methods and DL architectures.

## Introduction

Dynamic functional connectivity (dFC) is the variation of brain region functional connections over time derived from the Blood Oxygenation Level Dependent (BOLD) signals of regions extracted from functional magnetic resonance imaging (fMRI) data. dFC has been a growing discipline of research due to its importance in understanding brain processes and potential applications as a clinical biomarker for predicting the onset or tracking the progression of neurological diseases, including Parkinson’s disease (Engels et al., 2018), Alzheimer’s disease (Gu et al., 2020), and schizophrenia (Cattarinussi et al., 2023). dFC using task-based fMRI data examines the time-varying interactions between brain regions of subjects that were instructed to perform a cognitive task, in contrast to resting-state dFC. Previous studies have demonstrated the ability of short-term fluctuations in whole-brain functional connectivity patterns to classify scans according to task paradigms, highlighting the potential of dFC patterns to reveal insights into system-level behaviors of the human brain and their correlation with mental states (González-Castillo et al., 2015).

Recent studies have demonstrated that deep learning models are effective at leveraging the features of dFC to predict or distinguish subject traits, clinical disease status, and consciousness state. For example, a CNN (Convolutional Neural Network)-LSTM (Long Short-Term Memory) hybrid has been used to classify gender, achieving a high accuracy, and predict intelligence (Fan et al., 2020), while graph convolutional networks (GCNs) have shown strong performance in distinguishing Alzheimer’s disease from controls (An et al., 2020). Furthermore, Gomez et al. found that deep learning models can reliably distinguish between different states of consciousness, grouped by varying levels of awareness or arousal, by capturing dynamic patterns of whole-brain resting-state functional connectivity (2024). While there have been studies exploring the use of deep learning (DL) models on dFC in the last five years, the focus has been on using resting-state dFC to predict individual traits, disease status, or states of consciousness. Little work has been conducted to investigate whether task-based fMRI could provide robust dFC features for DL models to predict ongoing transitions in cognition. On the other hand, many statistical methods have been developed for dFC estimation, but the choice of dFC assessment methodology has been found to significantly impact the results of resting-state dFC, questioning the reliability and interpretation of such estimation techniques when applied in dFC studies (Torabi et al., 2024). In this study, we used dFC assessed by three methodologies, each of which captures a different aspect of the functional variations in cognitive activity, so that the analysis framework is more comprehensive and the downstream results are more reliable.

Our core investigation focuses on understanding how deep learning (DL) architectures can be developed and adapted to reliably predict task presence over time from task-based dFC time series and identifying which dFC assessment methods yield the most robust features for this supervised classification. In other words, the objective is to design and compare DL models for classifying whether a subject was either engaged in any instructed task or at rest, based on the corresponding dFC patterns at each time point. If the dFC features from a given dFC assessment method more consistently predict task presence, then it is more likely that dFC obtained by that assessment method is more representative of the latent functional connectome variations than other methods and consequently, should be incorporated into the dFC estimation pipeline of future studies. Hence, this work also establishes a framework to use DL prediction capacity to validate the robustness of select dFC assessment methods in capturing task-driven dynamics. To predict task presence, we developed a convolutional neural network (CNN), a node-level classification graph convolutional network (GCN), and a graph-level classification GCN. These models form the analysis pipeline of dFC assessed from an open-source multi-task paradigm, fMRI dataset from the OpenNeuro repository (Braver et al., 2025). There was also strong interest in how prediction performances vary across diverse task paradigms, which are the different types of cognitive tasks that the experimenters designed, so performances across task paradigms and across assessment methods were examined to identity latent patterns. To leverage the potential of DL, models were implemented by refining standard DL architectures to better represent and extract the unique characteristics of dFC.

## Methods

### Data, Setup, and Preprocessing

The data analyzed in this investigation is task-based dFC preprocessed from a task-based fMRI dataset from the OpenNeuro repository consisting of 38 subjects completing four task paradigms: AX-CPT (Axcpt), Cued Task-Switching (Cuedts), Sternberg (Stern), and Stroop collected by Braver et al. (2025). Each paradigm consists of two scans, and each scan has around 1081 time points and 100 brain regions of interest (ROIs), parcellated according to the Schaefer 100 parcel atlas (Schaefer 2018 parcellation, 2018). For clarity, we will consistently refer to the various types of cognitive tasks that a subject completed as task paradigms, whereas the time points when a subject is actively doing any task paradigm instead of resting as task.

First, to better understand the semi-structured dFC data and its characteristics as well as to formulate the problem setup, we provide some context on how BOLD signal time series were processed to form time series of dFC matrices. We binarized the task presence label (0 for rest and 1 for task) for each time region (TR) by removing TRs that are determined to be transition periods between rest and task states. Next, each truncated BOLD time series is assessed by three well-known dFC assessment methodologies: Time-Frequency (TF), Sliding Window (SW), and Coactivation Pattern (CAP) according to (Torabi et al., 2024). In general, these dFC assessment methods are statistical frameworks that aim to estimate the pairwise neural connection strengths between brain regions using correlation-based measures. For instance, sliding window is a standard method that involves calculating the Pearson’s correlation coefficient r_ij_ within a moving window of a fixed window length W, creating a time series of correlation coefficients representing the functional connectivity between *i* and j dynamically over t. Specifically, for a timepoint t, the correlation is computed on the window 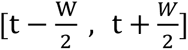 in the BOLD time series, and then the window is shifted horizontally to the next position in time. dF*C*_*ij*_ at time point *t* is defined as:

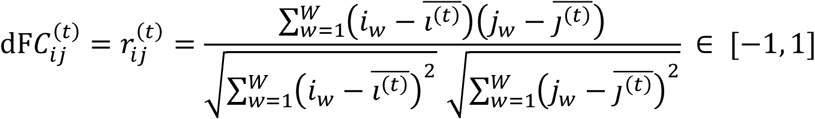

where 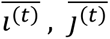 are the means of ROIs *i*’s and *j*’s signals, respectively, within the window for time point *t*, and *w* indexes over the TRs in each window. The higher the alignment between the two ROIs’ BOLD time series, the higher the correlation, and therefore, the more functionally connected those two brain regions are.

After dFC estimation of BOLD time series, we obtain dFC in matrix representation *M*_*t*_ ∈ ℝ^100×100^ each of size 100 ROI by 100 ROI, where each *m*_*ij*_ entry of 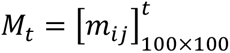 represents the strength of functional connectivity between the *i*th ROI and *j*th ROI, at a time point *t*. The dimensions of dFC matrix objects *M* for a given task paradigm *p* are the number of time points for a specific subject *T*_*s*_, and the number of subjects for that specific task paradigm *S*_*p*_. Additionally, *d* indexes over each of the 12 *dFC Assessment method* × *Task Paradigm* datasets. Each combination will be referred to as a unique (sub-)dataset in this study, and since we examine three dFC assessment methods, indexed by *a*, and four task paradigms, this produces twelve distinct datasets *D*_*d*_. Note that this, however, does not create a structured dFC tensor because each subject’s scan can have different numbers of time points, each dFC assessment method produces different numbers of time points *T*_*s*_ (e.g., sliding window yields the fewest number of time points for a fixed subject’s BOLD time series due to the window lengths), and not all subjects have participated in all four task paradigms. The semi-structured, tensor-like organization of dFC data constrains the analytical approaches that can be used. As a result, it is often need to down-sample specific matrices either across assessment methods or across the time series to impose a uniform tensor structure suitable for downstream analysis.

We consider and perform our prediction classifications on the dFC matrix time series M_t_, with a binary task presence label *y*_*t*_ ∈ {0, 1} at each time point, for each combination of task paradigm and assessment method dataset *D*_*d*_, individually. Therefore, the entire available dFC data *D* for the analysis is the union of *D*_*d*_, each of which are the unions of pairs of dFC matrix and its corresponding task presence label at each time point over all the subjects in that dataset indexed by s:

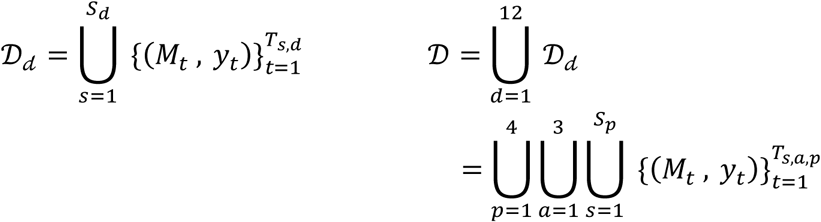

Next, a rank normalization was performed on each dFC matrix to ensure that the connectivity strengths are comparable across dFC assessment methods, since each method produces connectivity strengths on varying scales and of varying distributions. We note that each dFC matrix is symmetric since connectivity strengths are symmetric (i.e., the neural connectivity from the ith ROI to jth ROI is equal to the connectivity from the jth ROI to ith ROI), and the entries along the main diagonal are irrelevant since they represent each ROI’s self-connectivity, which is always the highest strength possible. Given these characteristics of a dFC matrix, only entries 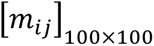 in its upper triangle (or lower triangle) are informative, and the number of these features is 100 ROIs choose 2, i.e., 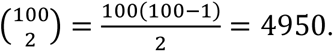 Then, the absolute values (magnitudes) of these distinct dFC features across all *T*_*s*,*a*,*p*_ dFC time points for one subject *s* at a specific dFC assessment method *a* and task paradigm *p* were ranked together. Therefore, raw dFC entries for all the matrices of one subject are now mapped by a bijection to m_ij_ ∈ {1, 2, …, 4950 ⋅ *T*_s,a,p_}. This procedure is repeated separately for each subject, assessment method, and task paradigm.

These steps were completed using the toolboxes fMRIPrep (Esteban et al., 2019), nilearn (Nilearn Contributors, 2023), and PyDFC (Torabi et al., 2024). For more details on the fMRI data processing pipeline to obtain dFC data, please refer to (Torabi et al., 2024).

### Deep Learning Models Implementations

We developed three deep learning architectures adapted for learning dFC patterns in Python: a CNN, a Node-Level GCN, as well as a Graph-Level GCN (based on the level at which time points are classified as rest or task). Deep learning pipelines were implemented using PyTorch (Paszke et al., 2019) and specifically the GCN framework was built using PyTorch Geometric (Fey & Lenssen, 2019). Data preprocessing and feature extraction in the exploratory phase were completed using Scikit-learn (Pedregosa et al., 2011). We ran all DL experiments systematically using McGill University, Department of Mathematics and Statistics’ GPU clusters: Nvidia H100 (96GB VRAM, 32GB RAM) and Tesla P100-PCIE (16GB VRAM, 32GB RAM).

For both training and testing, we implemented a cross-validation framework with five outer folds for reporting reliable performance estimates and three inner folds for hyperparameter tuning. This means that for each dataset, first it undergoes a randomized 80/20 train/test split grouped by subjects, then among the train samples, it undergoes a randomized 66/33 inner train/validation split grouped by subjects, resulting in approximately a 53/27/20 train/validation/test split.

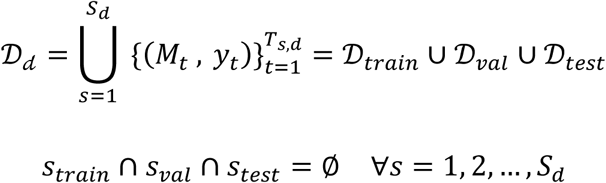

We used the Adam (Adaptive Moment Estimation) optimizer (Kingma & Ba, 2015) and its regularization version, AdamW (Weighted Adam) optimizer (Loshchilov & Hutter, 2017). For experiments that use AdamW, an L2 weight decay by default is directly applied to the model’s parameters and tuned over *weight decay* ∈ {0.0001, 0.001}. After the models learn parameters through hidden layers, the sigmoid function, 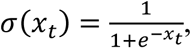 was applied to the scalar output of the linear layer *x*_*t*_. This is followed by a calculation of losses using the Binary Cross-Entropy with Logits Loss loss function, adjusted for the class imbalance between the number of samples with rest labels *N*_*neg*_ compared to those with task labels *N*_*pos*_ :

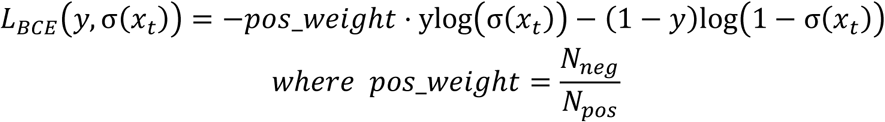

Lastly, for binary classification, a threshold function, 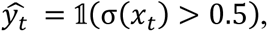 was applied to predict the task presence 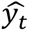 at time point *t* for all models and datasets.

### Convolutional Neural Network (CNN)

The CNN architecture was initialized from EfficientNet-B0, a widely used baseline CNN model that was pretrained on over a million images from the ImageNet database and achieves high accuracy with relatively few parameters (Tan & Le, 2019). It employs compound scaling to balance model depth, width, and input resolution. It is composed of several mobile inverted bottleneck convolution (MBConv) blocks with three by three and five by five convolutional kernels, which reduce computational complexity while preserving the learned feature representations. The basic convolution operation used in the initial stem and final head layers with *C* input channels, *K* output channels for 2D inputs 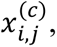 and kernels *W*^(*k*,*c*)^ of size *M* × *N* is defined as a cross correlation:

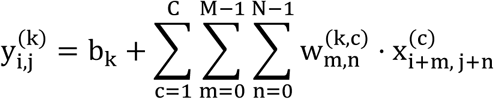

where 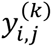 is the output of the feature map at the (*i*, *j*) position, *b*_*k*_ is the bias for output channel *k*, and 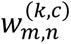 is the kernel weight at position (*m*, *n*) of kernel *W*^(*k*,*c*)^ that connects input channel *c* to output channel *k*. It also uses the Swish activation function SiLU(*x*) = xσ(*x*) between its hidden layers. This is a form of transfer learning since we then fine-tuned the CNN model to best fit and predict on dFC data. The main benefit is the reduced need for training on a large number dFC data samples across a large cohort size to reach the same level of performance because deep learning models are notoriously data hungry.

The EfficientNet-B0 architecture requires input samples to be of dimensions (224, 224, 3) where the last dimension represents the three channels of an RGB-colored image. For CNN input image construction, one way was to triplicate the values of each dFC matrix and stack them into a dFC tensor of shape (100, 100, 3), mimicking the RGB image channel’s structure. An alternative, multichannel way is to stack the three dFC matrices that originated from the same BOLD time series segment but were each assessed by a different dFC assessment method along a third axis to produce a tensor of dimensions (100, 100, 3). Again, each subject’s scan usually has a different number of time points, and each dFC assessment method produces different numbers of time points from the BOLD time series. Due to this semi-structured dFC data, implementing the multichannel method requires careful harmonization of the three dFC matrices along the number of samples in each time series to avoid a mismatch where some of the samples only have one or two dFC matrices stacked as a tensor input to the CNN model. The harmonization procedure keeps only the intersection of unique (subject, time point) tuples across the three assessment methods:

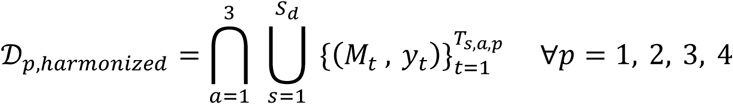

The choice of these two image construction methods was due to the natural integration of the square dFC matrices into the colored image format that the pretrained CNN model requires as input. Triplication is sufficient for turning dFC data into appropriate input data, but the multichannel way enriches a single sample’s information and was inspired by data augmentation approaches. For both ways, we transformed each dFC tensor to match the dimensions (via padding and resizing) and distributions (via transforming the per-channel means and standard deviations) of the images that EfficientNet-B0 was trained on before inputting them into our CNN implementation to improve the dFC data’s generalizability and transferability during training. For more details on the input transformations and deep learning model architecture, please see Fig. 2.

**Fig. 1:**
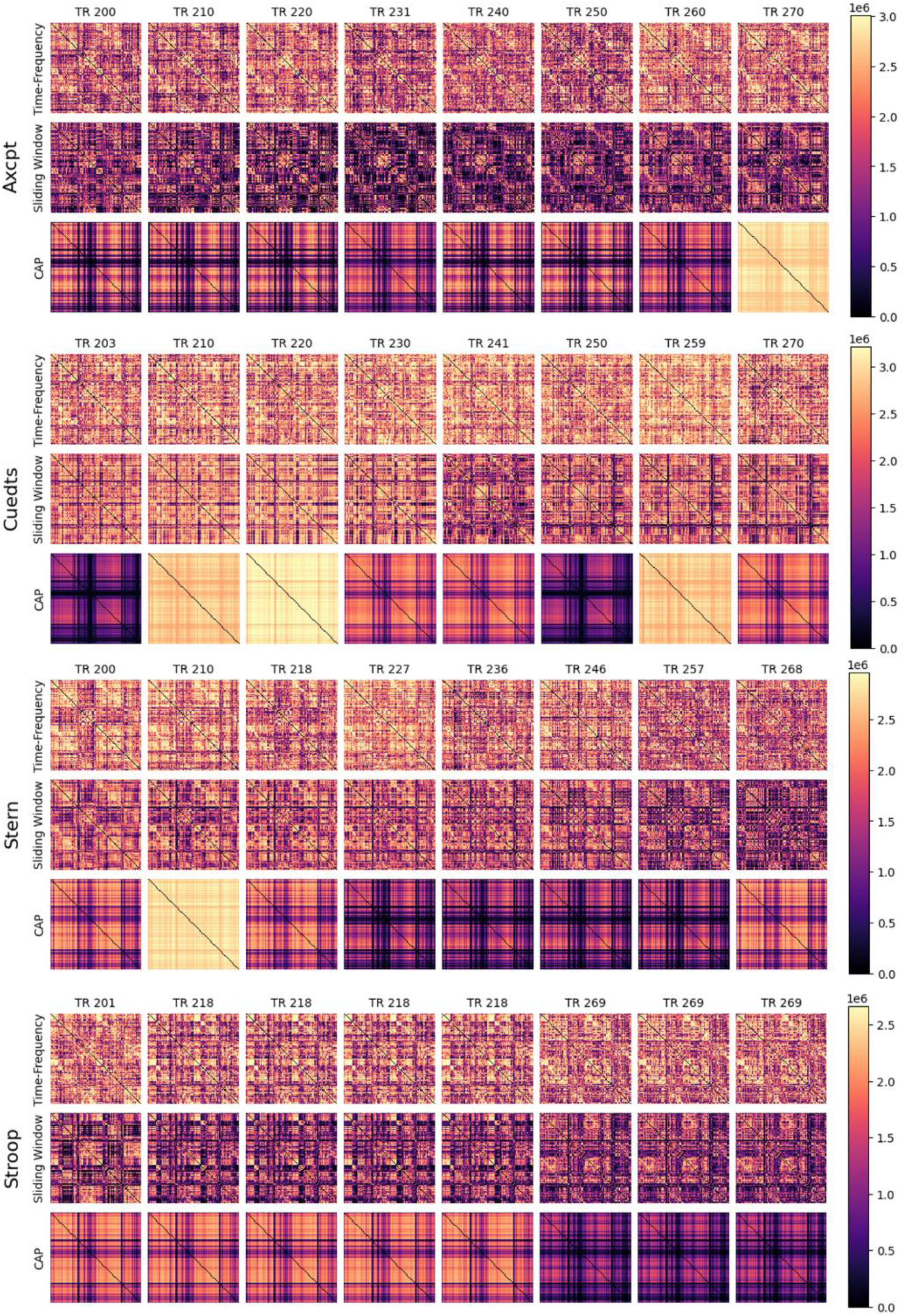
dFC data for one subject at select time regions (TRs) across all dFC assessment methods and task paradigms. The sequence of desired TRs was {200, 210, 220, 230, 240, 250, 260, 270} but many of those TRs are not available (either removed during the binarization of task presence or the harmonization of TRs across assessment methods). Thus, the calculated closest TR to each of the desired TRs are displayed, for each task paradigm. CAP is a method with a finite number of states, whereas the time-frequency and sliding window methods are state-free methods with an unlimited number of unique configurations evolving over time, seen in the figure.

**Fig. 2:**
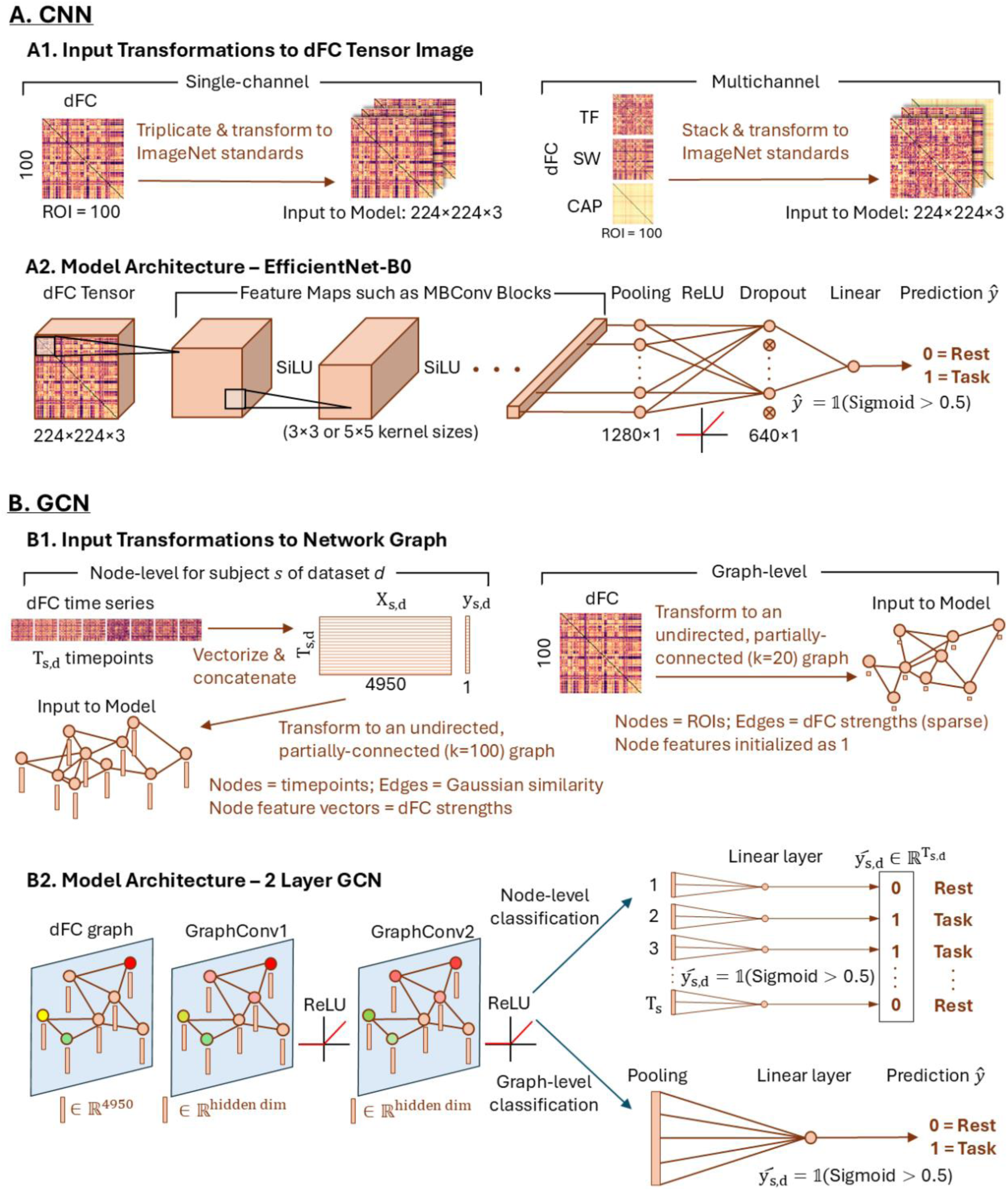
Deep learning dFC input transformations and models’ architectures. **A1.** Transformations of dFC data for single- and multi-channel CNNs. We transformed the dFC matrices to match the shape and distribution of images that the base model was originally trained on. **A2.** The model architecture is fully shared between both channel options. Due to limited subjects in the dataset, a pretrained EfficientNet-B0 model was used, followed by (global average) pooling, ReLU, dropout, and a linear layer. **B1.** Two distinct GCN frameworks for input dFC transformations depending on the level at which classification is performed. Node-level (left): For each subject of a given dataset, their dFC time points are vectorized and converted to a network graph. Each node represents a time point, its corresponding dFC vector is its feature vector, and the edge weights are the Gaussian RBF kernel similarities between each pair of time points. Graph-level (right): Each node represents a ROI, and edge weights for each node are either the dFC strengths, if it is connected to one of the strongest twenty nodes, or zeroed. **B2.** The initial layers of the GCN architecture are shared between the two classification frameworks. Then, for node-level classification, each node’s feature vector collapses and transforms into a binary prediction. One forward pass of one subject’s graph gives the task presence predictions for all its time points. For graph-level classification, an additional pooling layer aggregates the node features from the 100 ROIs before transforming it into a single binary prediction for one time point.

The hyperparameters were: {*batch size* ∈ {16, 32}, *learning rate* ∈ {0.0001, 0.001}, *weight decay* ∈ {0.0001, 0.001}, *dropout* = 0.5, *epochs* = 10}. Initial experiments tuning on larger sets of values for each of these hyperparameters revealed a slight preference for these specific values, so these were kept for large-scale performance computations. Relatively smaller learning rates were chosen since the model was pretrained, so only fine-tuning is necessary. An additional dropout layer was added after pooling and ReLU to enable the remaining neurons to learn more robust features and increase the model’s generalizability to unseen test data. In Fig. 5A, one can observe the overfitting and generalization challenge of this stochastic model, suggesting that only small numbers of epochs were needed to achieve the same or even better performance on test data and reduce training time (essentially an early stopping approach). This finding motivated why the reported results were for the model trained only on ten epochs, and this limitation will be addressed in the Discussion section.

### Node-Level Graph Convolutional Network (GCN)

Both GCN classification frameworks share certain dFC preprocessing steps and hidden layers. Before the downstream analysis, the entries of all the raw dFC matrices *M*_*t*_ were min-max normalized to [0, 1] by: 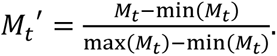 Then, we implemented two frameworks of a graph convolutional network based on the level at which the classifications are performed: node-level GCN and graph-level GCN. They were built from scratch instead of using a pretrained model, and the initial layers of their respective architectures are shared, alternating two times between a graph convolutional (GraphConv) layer and a ReLU (Rectified Linear Unit) activation function, ReLU(*x*) = max(0, *x*), applied on each node’s embedding vector. GraphConv encodes message passing between nodes and neighborhood aggregation for updating node feature embedding vectors, and it is followed by ReLU, which introduces nonlinearly and enhances the expressiveness of the network to learn complex patterns. This composition is defined by:

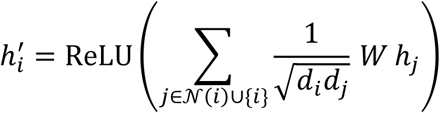

where 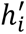 is the updated embedding vector of node *i*, *N*(*i*) is the set of node *i*’s neighbors, *d*_*i*_ is the degree of node *i*, *W* is the learnable weight matrix between nodes, and *h*_*j*_ is the embedding vector of neighboring node *j*. Then, the GCN model’s architectures diverge for performing a node-level or graph-level classification of task presence. For more details, please see Fig. 2.

Specifically for the node-level GCN model, we predict task presence at each node. For its input dFC transformations, for a fixed *dFC Assessment Method* × *Task Paradigm* dataset *D*_*d*_, we create an individual graph for each subject *s*. For each of time point in that subject’s time series data, the upper triangle of its symmetric dFC matrix is vectorized to *x*_*t*_:

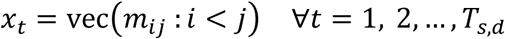

These dFC vectors and their corresponding true labels 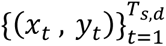 are concatenated row-wise to form matrix 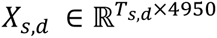 and labels vector 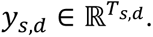 Next, dFC data was converted to a network graph for each subject. Each node represents a time point *t* where its corresponding dFC time point vector *x*_*t*_ is its feature vector, and the edges are initialized as the L2 Gaussian RBF (Radial Basis Function) kernel similarities between pairs of time points with σ = 1:

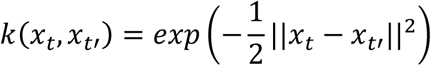

The hyperparameters for this model were: {*k* = 100, *sigma* = 1, *learning rate* ∈ {0.0005, 0.001}, *hidden dim* ∈ {16, 32}, *weight decay* ∈ {0.0001, 0.001}, *epochs* = 300}. After exploring a sparse but broad range of values for *k* and *sigma*, we found that 100 neighbors are sufficient to produce graphs that balance local and global connections well, and the value for sigma is not important, so it is set to 1 to subsequent runs. Hidden dimension is a hyperparameter that specifies the number of features for each node’s embedding vector to keep after each GraphConv layer. It was found that for the Node-Level GCN, prediction capacity scales with the number of epochs up to around 300 epochs, unlike the other two models implemented in this study, so the model was trained over a larger number of epochs for each fold, and this observation will be investigated upon in future work. Graphs were not batched when inputting them into the model because each of them represents one subject, so there is a maximum of only 38 graphs for any dataset. For a fixed dataset, one forward pass of one subject’s graph through the node-level classification model yields the task presence predictions for all the time points of that subject’s scan.

### Graph-Level Graph Convolutional Network (GCN)

Aside from the shared GCN components, the graph-level GCN model has a distinct setup because its goal is to predict task presence for each graph. First, we transform each dFC time point’s matrix to an individual graph, where each node in the graph corresponds to a brain ROI, all the node features were initialized to a constant, and each min-max normalized dFC strength between ROIs *i* and *j* was implemented as the edge weight in the graph, i.e., 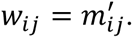 This graph construction seems natural, mimicking the functional organization in the brain where the brain signals originated from. As an aside, three graph edge options were explored to make the graph sparser: fully connected (leave the graph as is), *k* nearest neighbors (only keep the directed edges from each node to the *k* nodes whose edge weights are the highest), and thresholding (only keep edges whose weight is at least 0.5). However, each option after optimizing over its unique hyperparameter did not lead to significant classification performance differences, so the default graph setup used *k* nearest neighbors.

The hyperparameters for this model were: {*k* = 20, *batch size* ∈ {16, 32}, *learning rate* ∈ {0.0005, 0.001}, *hidden dim* ∈ {16, 32}, *weight decay* ∈ {0.0001, 0.001}, *epochs* = 10}. Since there are a total of 100 nodes in each graph, keeping only 20 neighbors per node was adequate to preserve local and some global structures. For graph-level classification, an additional pooling layer is needed to aggregate the node features from the 100 nodes because there is only one task presence label per dFC time point graph. We used global mean pooling to get the graph-level embedding *h*_*t*_ at time point *t* defined as: 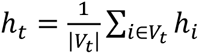 where *V*_*t*_ is the set of all nodes in the graph at time point *t*, and *h*_*i*_ is node *i*’s embedding vector after being updated by the last ReLU(GraphConv) hidden layer composition. Finally, a linear layer and transformations output a single binary task presence prediction per graph for comparison with the true label.

## Results

The core goal of this study is to understand how deep learning (DL) model architectures can be adapted to predict subjects’ binary task presence over time utilizing their task-based dynamic functional connectivity (dFC) data and to identify which dFC assessment methods yield the most robust features for classifying time points. The test performances were evaluated using two metrics, balanced accuracy (accuracy adjusted for class imbalance) and Area Under the ROC Curve (AUC), each defined as:

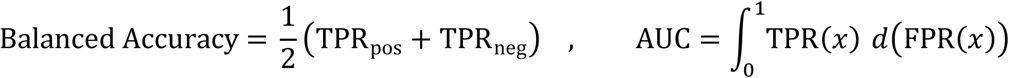

where 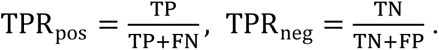 Both metrics have 0.5 as the chance-level performance, the performance attained if the machine guessed the binary task presence label randomly. From Fig. 3A, the CNN and Node-Level GCN implementations consistently outperform the current implementation of the Graph-Level GCN across all the datasets analyzed. However, this is not because the CNN models are effective, in fact their average balanced accuracies on all combinations of task paradigm and assessment method are below 75%, but because the Graph-Level GCN is insufficient to handle the complexity of the noisy data. For CNN, the multichannel method’s balanced accuracy is within the performance range of the other three methods for all the task paradigms, which could be because it was constructed as a hybrid of the other three, and this observation is further discussed in the Discussion. The Graph-Level GCN model consistently displays chance-level performance, with small standard deviations across folds, suggesting that it has trouble learning any patterns in the data. Examining Fig. 3B, after aggregating each model’s performances by dFC assessment method, it is evident that for both CNN and Node-Level GCN, Time-Frequency shows the best performance, CAP shows the worst performance, and sliding window is the most variable. Sliding window’s large error bars overlap with the error bars of all the other assessment methods for CNN, and its performance is increased when using the Node-Level GCN model, compared to the CNN, despite the smaller input sample size. After grouping by task paradigm, we notice that for all three DL models, their performances across the four paradigms do not have a significant difference from one another since the interquartile ranges all show significant overlap.

**Fig. 3:**
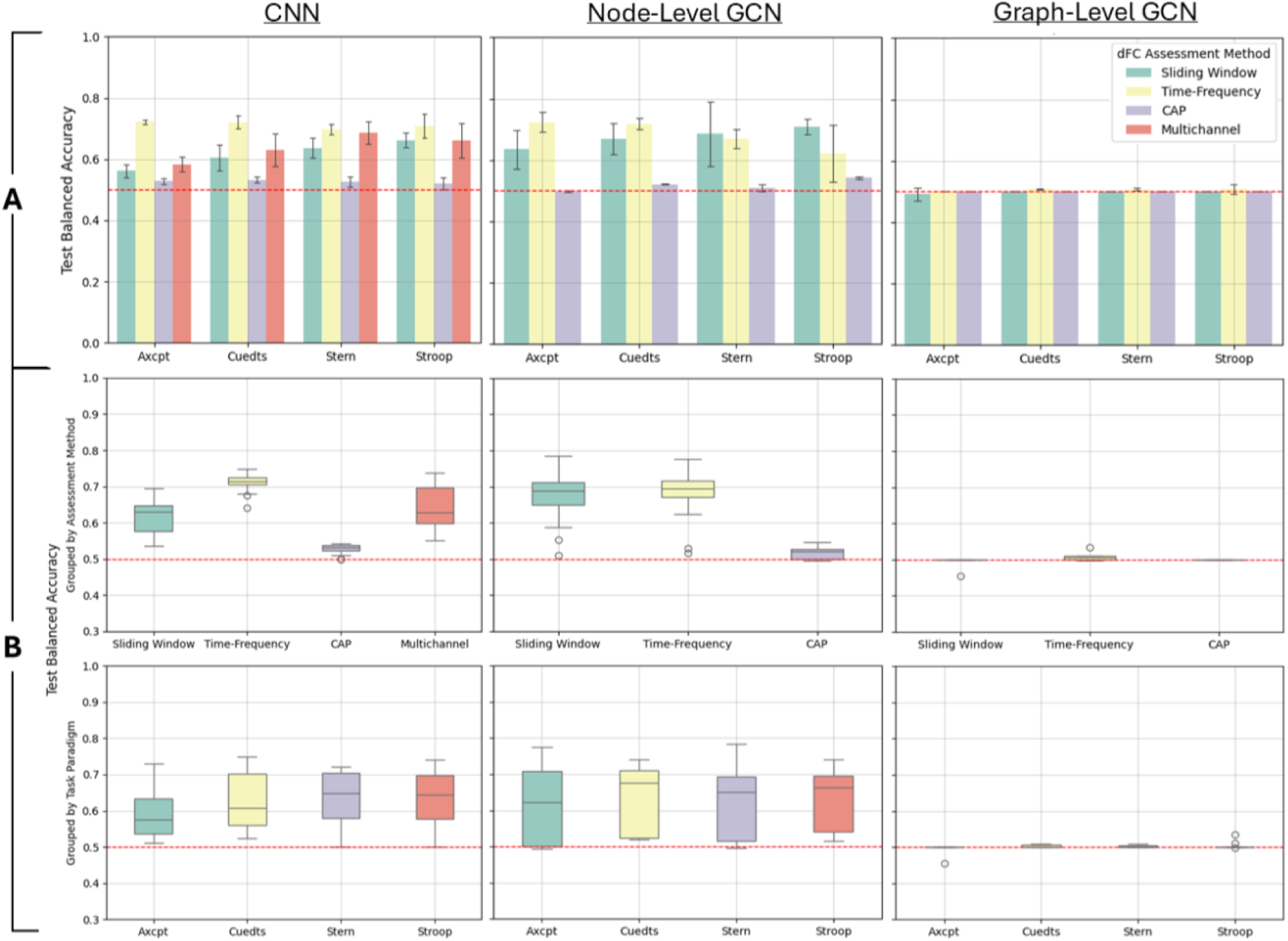
CNN, Node-Level GCN, and Graph-Level GCN balanced accuracy test performances for all datasets, averaged across five cross validation folds. The dotted red horizontal lines indicate chance-level performance. **A.** Individual *dFC Assessment Method* × *Task Paradigm* dataset performances. The multichannel method, which combines data from the three dFC assessment methods, is only an option for CNN. Error bars indicate ± 1 standard deviation across five folds. **B.** Summary of test accuracies from A, grouped by assessment method (row 1) and by task paradigm (row 2). Due to the small deviation of Graph-Level GCN performances away from 0.5, implying small interquartile ranges, several performances are illustrated as outliers when they are simply showing consistent poor performance.

After comprehensively running experiments for all three DL models (CNN, Node-Level GCN, and Graph-Level GCN) on all twelve sub-datasets *D*_*d*_, we can interpret from Fig. 3 that both the choice of DL model and dFC assessment method significantly influence the prediction performance on the test dataset time points, while the task paradigm that the subject was performing during data acquisition showed minimal impact on this task presence classification performance. Fig. 4 shows the same test performances for each combination of DL model and sub-dataset but visually as a performance heatmap with specific scores, as evaluated by both balanced accuracy and AUC metrics. AUC displays the same trend as balanced accuracy in which the Graph-Level GCN model and the CAP assessment method show the worst performances when either one is crossed with any other valid option, while Time-Frequency seems promising for any task paradigm when analyzed by a CNN or Node-Level GCN model.

**Fig. 4:**
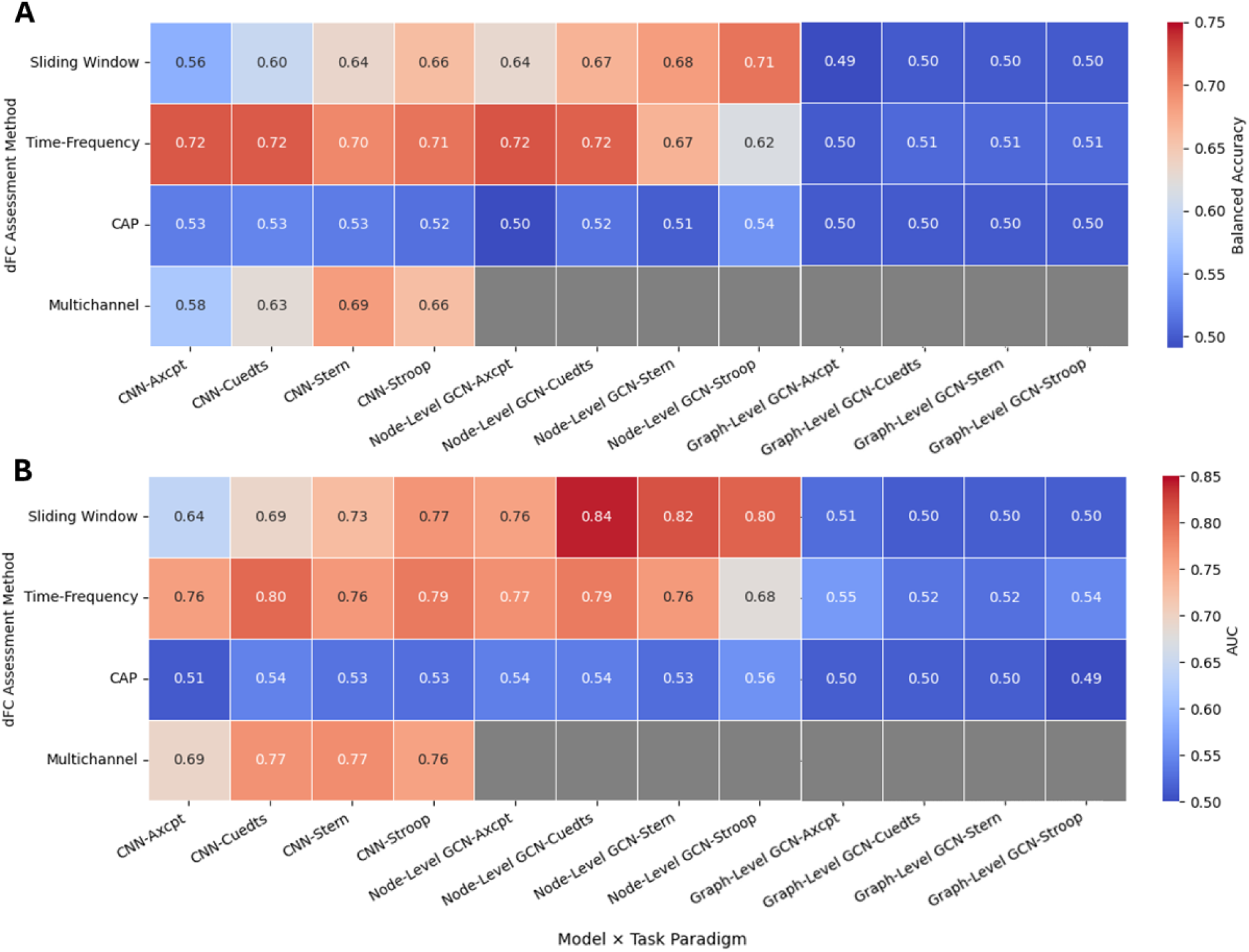
Heatmap of numerical test performances for all models on all datasets, averaged across cross validation folds. **A.** Balanced accuracy scores. **B.** AUC scores. Chance-level performance is 0.5 for both metrics. Multichannel is only an option for CNN, so GCNs × Multichannel boxes have no scores.

To investigate the training behavior of models and the extent to which regularization and early stopping approaches should be applied, we examined the models’ learning curves across epochs in Fig. 5. In subplot A of Fig. 5 showing the CNN model, for the evaluation (validation and test) losses, there appears to be an increasing trend of losses that scales up linearly with epochs. There is no intersection between the training and test curves, contrary to most DL learning curves, and test losses are always significantly higher than train losses, indicating that the current implementation of the model is overfitting prematurely. Therefore, as training time increases, the model performs worse on test data, and this was the motivation for why in the performance metrics reported above, we opted to run experiments with only ten epochs. In subplot B, noticing the small y-axis scale, all the losses are stagnant over epochs. Not even training loss decreases, suggesting that the Graph-Level GCN model is underfitting and cannot fit well to the dFC data characteristics in general. This may explain the chance-level performance across all datasets. In subplot C, there are huge peaks in the Node-Level GCN’s training and test losses that occur approximately in phase with each other and periodically every twenty epochs.

**Fig. 5:**
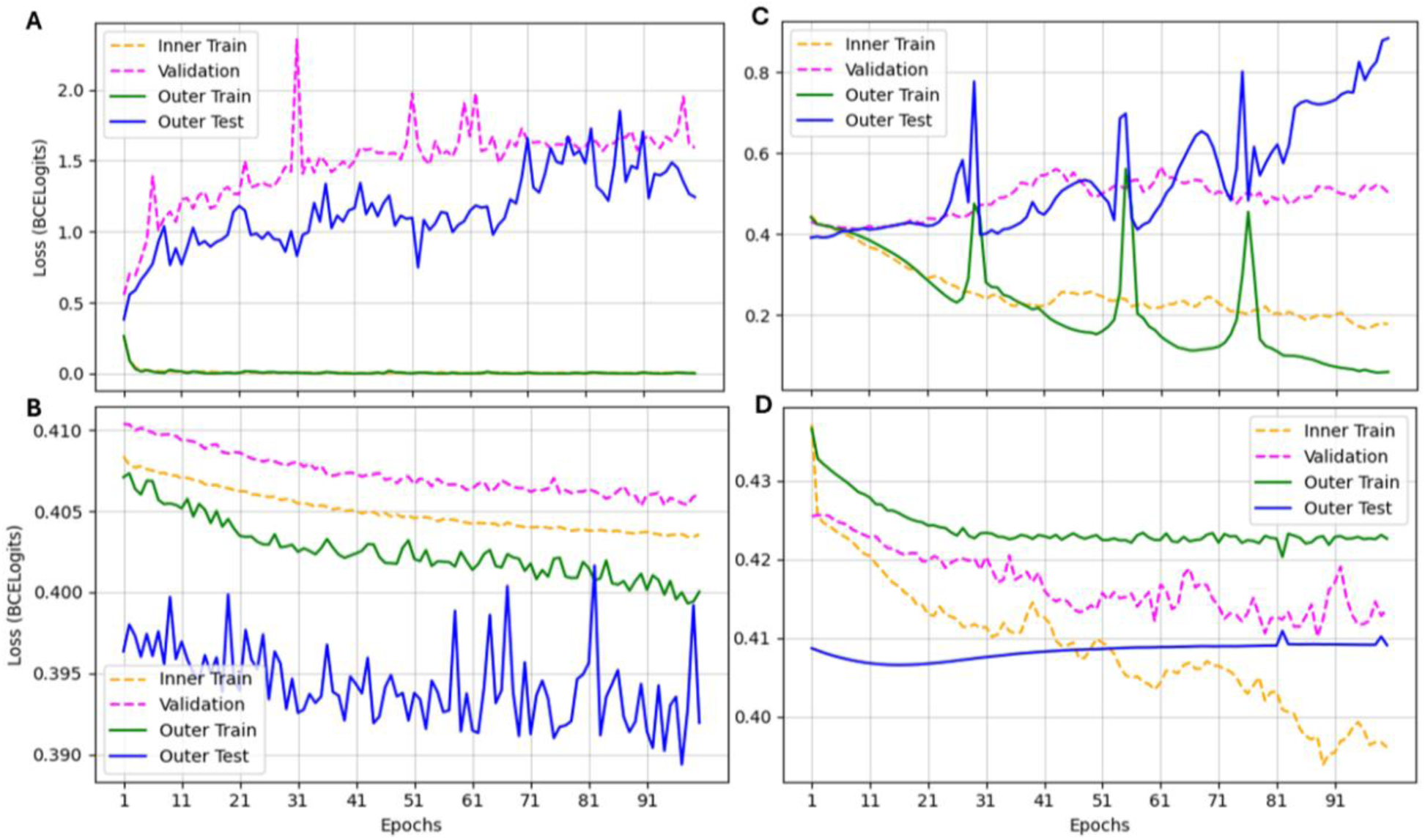
Binary Cross-Entropy losses after one to 100 epochs for various DL models. Depicted are the learning curves for fold one, trained and evaluated on data from the Axcpt task paradigm assessed by the Time-Frequency method. The sub-plotted models are **A.** CNN **B.** Graph-Level GCN **C.** Node-Level GCN **D.** Heavily regularized Node-Level GCN with additional *dropout* = 0.5 applied after each graph convolutional layer. The inner train curve shows the losses averaged over all combinations of hyperparameters and inner folds, and the validation curve shows the corresponding averaged losses on the reserved validation sets. The outer train and outer test curves show the performance of models trained and evaluated on the entire dataset with the tuned hyperparameters.

This trend is largely due to the stochasticity of the Adam optimizer and is likely aggravated by the small subject cohort size that is equivalent to having a small sample size in this model. In subplot D, we show an example of applying heavy regularization penalties to the Node-level GCN. This experiment added additional regularization (*dropout* = 0.5) after each ReLU(GraphConv) composition layer to decrease the overfitting behavior seen in C. However, this caused the training loss to plateau across epochs and introduced a trade-off in test performance. Specifically, this regularization approach resulted in roughly a 10% drop to a balanced accuracy of 0.62 and an AUC of 0.67, ran using the same hyperparameters as in other node-level GCN experiments. This provides evidence for why models with heavier regularization procedures were not used in running experiments comprehensively over all datasets, despite observing substantial overfitting behavior in the CNN and Node-Level GCN models.

At the current stage, we found that the factors that influence test performance, in order of descending importance, are DL model, dFC assessment method, and task paradigm. All datasets trained and evaluated using the CNN and Node-Level GCN implementations demonstrate better task presence predictive capacity than all datasets trained and evaluated using the current Graph-Level GCN architecture. However, the performances of CNN and Node-Level GCN are similar, with the exception that the performance of Node-Level GCN applied to dFC data assessed using the sliding window method is elevated from that of CNN by 8-10%. Then, given that the model is not the Graph-Level GCN, datasets assessed by the CAP method consistently underperform compared to the other dFC estimation methods. Performance differences are very similar across the four task paradigms.

## Discussion

We observed that the prediction performances were primarily driven by which of the three deep learning (DL) architectures were applied to train and evaluate the dynamic functional connectivity (dFC) data, and secondly by which of the three dFC assessment methods explored in this study were used to estimate dFC from Blood Oxygenation Level Dependent (BOLD) signals from fMRI scans. This result highlights the issue of excessive analytical flexibility in both the fMRI to dFC processing pipeline via dFC assessment methods, and more fundamental to this study, the dFC to task presence classification pipeline via DL models. Moreover, because the task paradigm that subjects were performing during data acquisition had minimal influence on prediction performance, these results suggest that methodological choices may have a greater impact on dFC-based predictions than the neurocognitive factors themselves. This raises the concern that analysis methodology may overshadow cognitively meaningful variables, complicating the interpretation of cognition state predictions using dFC features.

Our results indicate that DL models can detect latent differences in brain dFC patterns and structures between when a subject was cognitively at rest or doing a task, for all the task paradigms analyzed. However, this is primarily dependent on the DL architecture used on the dFC data for the task-presence predictions since the CNN and Node-Level GCN both noticeably improve the prediction accuracies of time points from subjects that were unseen during training, while the Graph-Level GCN fails to improve the prediction beyond chance-level guessing.

Moreover, it was found in Fig. 4 that for all task paradigms, models predicting task presence on dFC assessed using the Co-activation Patterns (CAP) method yielded suboptimal, close to chance-level performances. This serves as evidence that CAP may not be the most effective method to estimate dFC from BOLD fMRI signals in this context since its downstream dFC data is significantly less predictive of the cognition state transitions than the other two dFC assessment methods examined in this study. However, a critical limitation of this investigation is that only four task paradigms were examined, so the claim that task paradigm has insignificant impact on cognition prediction over time should be interpreted with caution for other task paradigms. Another limitation, especially for the Node-Level GCN results, is that since each graph is constructed for an individual subject and there are a total of 38 unique subjects in the task-based fMRI dataset we analyzed, the total number of samples passed through the model for each epoch is at a maximum 38 graphs (since not all subjects participated in all four task paradigms), which is not sufficient for data-hungry DL models. Future work can perform the analysis on a more diverse range of cognitive task paradigms to see whether they would have varying prediction rates of task presence and on larger datasets with more subjects to improve the robustness and predictive performance of the models. Another idea could be to formulate the objective as multiclass classifications where instead of the prediction being whether a subject was performing any cognitive task other than rest at a given time point, the goal would be to predict which specific cognitive task (or rest) a subject was performing over time after combining all the data across task paradigms to be inputted to each model.

Furthermore, an interesting observation is that the performance of the multichannel method in the CNN model is in between that of the three dFC assessment methods that it was constructed from, for all the task paradigms. This raises the question of why the multichannel performance could not perform at least as well as the best performance of the three assessment methods. Such performance was expected since the best performing assessment method’s data matrices were directly supplied as one of the channels to the model during training. Although the channels are treated individually with separate convolution filters for certain layers, there are steps such as in the MBConv blocks in the EfficientNet-B0 model that perform aggregation of information across its input channels, so it is reasonable to question why the model did not learn to only or more heavily rely on the parameters given by the most useful dFC assessment method’s channel. This highlights a critical limitation of our work: the performance results lack interpretability in relation to cognition and physical brain regions. Moving forward, we aim to identify features that yield more robust signals of functional variations and understand why certain feature representations can improve the interpretability and performance of models.

Possible approaches include visualizing saliency maps to identify which input features contribute most to influencing the final prediction and labeling spatial brain region attributes such as three-dimensional physical location and hemisphere information to better understand if certain physical characteristics drive the downstream task presence prediction over time.

Regarding the GCN models, it is intriguing to compare their test performances and running times, as well as discuss future graphical extensions. First, we observe in Fig. 3 and Fig. 4 that the Graph-Level GCN consistently produces predictive performances closely distributed around the chance-level performance. This serves as potential evidence that building a model that locally models each dFC time point as a graph with brain ROIs as nodes to mimic the physical brain functional connectome is not effective at driving the prediction of cognitive state transitions over time. Furthermore, running experiments using the same GPU cluster to compute the fold one performance of Graph-Level GCN compared to Node-Level GCN, both with same hyperparameters and on the same Axcpt × Time-Frequency dataset after the same number of epochs revealed that these architectures also have running times that differ by orders of magnitude. The node-level classification setup is much more computationally efficient, having a running time of five minutes compared to the graph-level classification model’s running time of around twelve hours. A plausible explanation could be the substantially larger number of, albeit smaller, graphs that need to be batched when fed into the model for each epoch. The Node-Level GCN, which performs binary classification for each time point represented as a node in its subject’s graph, also opens the possibility for a hierarchical graph construction and model architecture that combines the Node-Level and Graph-Level GCNs. Specifically, it preserves the idea of performing a node-level classification where one creates an individual graph for each subject and each node is a distinct time point, but the features for each time point’s node becomes a secondary graph with ROIs as its nodes. Considering that the Node-Level GCN showed significantly better performance from this study, this hierarchical construction that is primarily structured in a similar way but additionally captures the physical functional connectome structure of the Graph-Level GCN could connect subjects, time points, and ROIs in a more cohesive and interpretable way. This could especially be insightful if we can encode meaningful spatial information about the ROIs into the edges of the subgraphs instead of using constant initializations for edge weights as in the current Graph-Level GCN implementation.

Additional interest lies in exploring the integration of DL models that were pretrained and designed initially for classifications on preferably dFC (or at least neuroimaging) data. The EfficientNet model was pretrained to perform classifications on, for instance, animal images from ImageNet, so it may be less effective at capturing the fine patterns of dFC matrices as tensor image inputs. In fact, viewing Fig. 1, it seems that the dFC matrices across time points all look very similar to the human eye regardless of them being the patterns at rest or at task, in contrast to the ease of distinguishing between whether an image depicts a cat or dog. Thus, the dFC task presence prediction objective is arguably also much more challenging to the DL model, especially when its model parameters were optimized for a drastically different classification.

From prior studies, Ngo et al. (2020) proposed BrainSurfCNN, while Madsen et al. (2024) proposed BrainSERF and BrainSurfGCN, all of which are deep learning models that were built to predict individual cognitive task contrasts, such as the brain activation maps between when a participant was solving a math versus a language problem, using subject’s resting-state functional connectivity data. These architectures could be promising and more transferable for enhancing this investigation which seeks to use task-based dynamic functional connectivity to predict binary task presence over time.

Although tuning models are not the focus of this investigation, a limitation of the reported results is that they are highly dependent on the choice of regularization strength for each model. In Fig. 5, it is expected to observe stochasticity in the loss learning curves because of the choice to use the Adam optimizer (with weight decay), a stochastic gradient descent method to update the models’ parameters. More notably, the bias-variance trade-off in choosing the appropriate regularization strength for the three different model architectures is evident in the prediction performances reported in Fig. 4 and the loss curves shown in Fig. 5. For the CNN and Node-Level GCN models (Fig. 5A, 5C) that achieve relatively higher prediction performances, the models have overfit significantly and prematurely to the training data and its noise since the training losses decrease while test losses increase over epochs. This is due to having large variances across different data subsets, making the models lose generalizability on the dFC data of unseen subjects. On the contrary, for the Graph-Level GCN and more heavily regularized Node-Level GCN models (Fig. 5B, 5D), one can observe that the models are underfitting since train losses are stuck at high losses as epochs increase, due to models having large biases to the true, unknown data distribution. Aside from noting that a future step could be to more extensively tune over a dense grid for hyperparameters that better balance this trade-off, it is interesting that for the overfitting CNN and Node-Level GCN models, both of their test prediction performances decreased in experiments with increased regularization penalties and approaches. Various additional regularization options, including weight decay with heavier L2 penalties, dropout after GCN layers, batch normalization, and freezing hidden layers of the pretrained CNN model, were explored but we found that either they still cannot effectively prevent the model from overfitting or they reduced overfitting seen in the loss curves at the expense of significant reduction in test prediction performances. Although it seems contradictory in definition, it is possible for test performances (balanced accuracies and AUCs) and test losses to simultaneously be higher in the overfitting scenario when the models are correctly classifying more time points but the few time points that they misclassified were misclassified with high confidence, leading to greater Binary Cross Entropy losses when compared with the true labels. Furthermore, the Node-Level GCN was found to have test performances that scale up as the number of epochs increases, despite overfitting, whereas the other two models suffer from decreased or stagnant metric performances over epochs. The underlying mechanism of these DL architectures driving these patterns of simultaneous overfitting and increased test prediction accuracies remains unclear and requires further investigation to understand. Future work includes implementing a basic, single-convolutional layer CNN and single graph-convolutional layer Node-Level GCN and experiment with applying them on both simulated and read dFC data to better understand the overfitting behaviour. This single hidden layer approach is supported by the Universal Approximation Theorem for feed-forward neural networks in theory, but we aim to test such simplistic models to predict on dFC data.

As an aside, there were tremendous intra-subject time point correlations in datasets across dFC assessment methods and task paradigms. This observation could be predominantly attributed to the dense number of time points (about 500-600 TRs per one fMRI scan of a subject after preprocessing) compared to the number of unique subjects. In the exploratory data analysis phase, we had observed that the UMAP biplot showed each subject’s dFC trajectory over time as a continuous, string-like curve that is separate from that of other subjects. When we ran our CNN model without grouping samples by subjects (i.e., using StratifiedKFold instead of StratifiedGroupKFold in doing the train/test splits in the inner and outer folds of cross validation), we found that the test performances were drastically higher, almost perfect predictions, than that after grouping, achieving a test balanced accuracy of 0.980 ± 0.007 and test AUC of 0.992 ± 0.009 on the Axcpt Time-Frequency dataset using the CNN. Despite this, our current, subject-grouped sample splitting procedure is fair because we should only be evaluating our model’s performance on the data of completely unseen subjects. Assigning all the time points of a subject into one of the train, validation, or test sets (mutually exclusive) is crucial to avoid subject-level data leakage. This result reinforces the challenge of generalizing results conducted for a sample of the population to the broader population, and the need for analyzing large and diverse datasets to capture population-level variability in the scientific findings. Moreover, this result emphasizes the importance of and potential applicability for rigorous statistical validation procedures and theoretical methods to reduce overfitting and improve the reliability of findings obtained from analyzing small sample sizes due to various constraints. In clinical applications, this finding may also motivate the design of adaptive experiments that can detect subgroups within a study, such as to account for subject heterogeneity in neural dynamics such as dFC, and enable personalized treatments, which is consistent with the goals of precision medicine.

This investigation has insightful findings in the interdisciplinary field of neuroinformatics. On the dFC estimation level, comparing task-presence prediction performances builds upon previous work that shed light on the methodological issue of analytical flexibility in the arbitrary selection of dFC assessment method, giving rise to the challenge of comparing results across applied dFC studies conducted by different research groups using different dFC assessment methods. On the task-presence prediction performance level, developing computational machine learning techniques that are adapted to the specific data type and properties of dFC matrices give insight into the drawback of analytical flexibility in designing and enhancing DL models by experimenting with an arbitrary architecture and set of hyperparameters. Thus, our work yields valuable knowledge for researchers when designing the analysis pipeline for their dFC research studies to be more robust and reliable. Overall, the investigation underscores the analytical prowess but also limitations of using DL techniques to derive interpretable findings from high-noise and high-dimensional neuroimaging data.

## Conclusion

The investigation demonstrated that deep learning models have strong potential to be adapted to predict task presence over time using task-based dynamic functional connectivity (dFC), while also shedding light on the significance of methodological choice in dFC estimation in biasing predictive performance. By exploring convolutional and graph-based neural network architectures, we underscored the importance of designing robust model implementations and dFC representations to execute stable and reliable classification. These findings illustrate the promise of deep learning to decode cognitive states from fluctuations in brain region connectivity signals over time. The results also urge the need for standardized practices and methodological transparency for advancing the use of dFC as a potential biomarker for task-driven brain state transitions in clinical settings. All data and code to produce the results and figures from this investigation are available upon request.

